# Trace gas oxidation supports sub-surface microbial communities across Namib Desert fog and aridity gradients

**DOI:** 10.64898/2026.02.19.706496

**Authors:** Dana Z. Tribbia, Pedro H. Lebre, Xabier Vázquez-Campos, Angelique E. Ray, Thomas Laird, Náthali Machado de Lima, Gillian Maggs-Kölling, Don A. Cowan, Belinda C. Ferrari

## Abstract

Widely accepted climate predictions indicate that drylands will expand to cover more than half of the Earth’s terrestrial surface by the end of the 21^st^ century. In these environments, harsh conditions including nutrient and water limitations restrict plant and animal life, thereby increasing the importance of soil microbial communities in nutrient cycling and ecosystem functioning. The Namib Desert is a distinctive dryland ecosystem characterised by a steep natural aridity gradient, transitioning from a coastal hyperarid zone influenced by frequent fog deposition to an inland arid region receiving seasonal rainfall. This study investigates the impact of water availability and moisture regime on microbial trace gas oxidation and community composition across this aridity gradient. Quantitative analyses revealed that total microbial abundance and activity indicators, including ATP concentrations and respiration rates, were significantly (*p* < 0.005) reduced in hyperarid soils compared to their arid counterparts. In contrast, hyperarid fog-dominated soils exhibited significantly (*p* < 0.0005) elevated rates of atmospheric hydrogen oxidation, even in the absence of water inputs. We propose that sustained high-affinity hydrogen oxidation, coupled with rapid microbial resuscitation following wetting events, supports shallow sub-surface microbial communities in the Namib Desert, particularly in the coastal hyperarid zone. Together, these findings challenge current understanding of the lower limits of microbial activity and reveal alternate metabolic pathways that enable microbial persistence in hyperarid hot desert soils.

**Importance:** Drylands are expanding globally, yet the mechanisms that allow microbial life to persist under extreme and sustained water limitation remain poorly understood. This study demonstrates that atmospheric trace gas oxidation, particularly high-affinity hydrogen oxidation, supports active and resilient microbial communities in hyperarid soils of the Namib Desert, even in the absence of liquid water inputs. By revealing how microbes may couple trace gas metabolism to energy and water generation, our findings provide new insight into the lower limits of microbial activity in dry hot desert soils and highlight the need to investigate how microbes persist and sustain soil ecosystem functioning.

## Introduction

### Water availability across a natural xeric gradient in the Namib Desert

The Namib Desert is a distinctive dryland ecosystem situated along a steep natural aridity gradient on the west coast of Namibia [1, 2]. Low intensity rainfall events occur intermittently (< 18 days/year) along this gradient [3, 4]. Precipitation frequency and volume increase progressively from west to east, with the coastal region receiving the lowest average annual rainfall (∼10 mm/year) [4]. These sporadic rainfall events are separated by prolonged dry periods, leaving much of the soil sub-surface desiccated for lengthy periods of time [3, 4]. The Aridity Index (AI), defined as the ratio of precipitation to potential evapotranspiration, divides the Namib Desert into two primary zones: a hyperarid coastal region (AI < 0.05) and a central arid region (0.05 ≤ AI < 0.2) [5, 6] (Figure 1A). Despite receiving extremely low annual rainfall, the hyperarid coastal zone experiences frequent fog events, approximately 3–9 days/month, which decrease in frequency and duration beyond 60 km inland [2, 4, 7].

**Figure 1:**
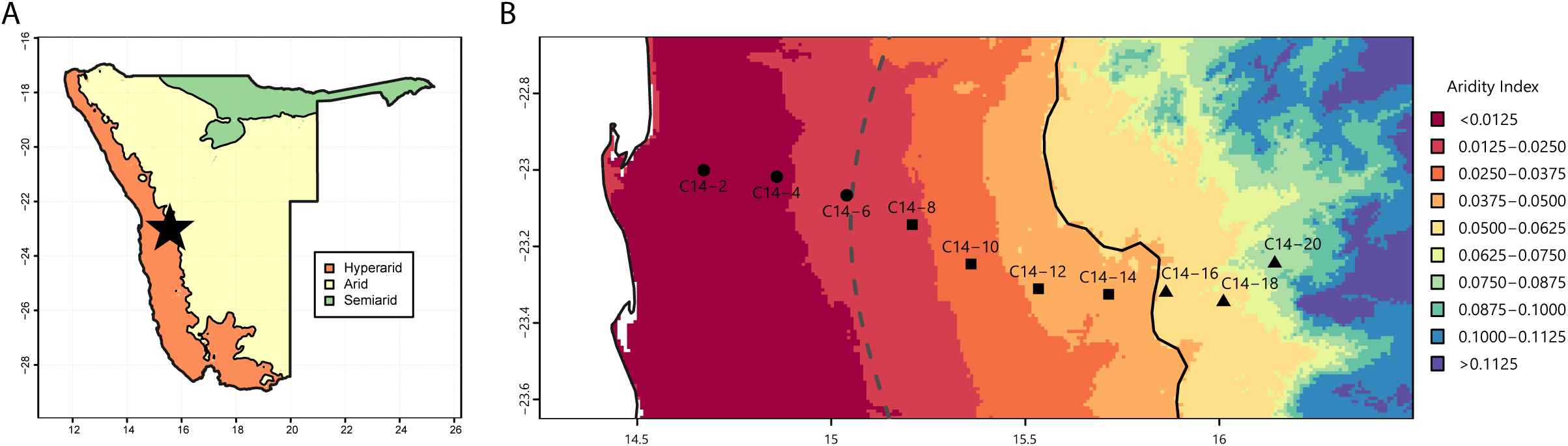
Aridity gradients of the Namib Desert and the C14 sampling transect. A) Xeric zonation of Namibia according to Aridity Index; transect location represented by the black star. B) GPS locations of ten sampling sites spaced 20 km apart along the 200 km C14 transect, located across a steep hyperarid-arid aridity gradient. The Fog zone is marked by the dotted grey line, and the hyperarid-arid boundary is marked by the solid black line. Fog, Hyperarid and Arid zone sites are indicated at circular, square and triangular points, respectively.

Considering the minimal precipitation along the hyperarid coast, fog-derived moisture has been proposed as the dominant source of bioavailable water for these soil microbial communities [3, 8–10]. Fog inputs are estimated to contribute up to 183 mm deposited moisture annually [2, 4, 7], and contribute to slight increases in sub-surface (0–5 cm) soil relative humidity, compared to soils from rainfall-dominated zones [3]. However, their limited capacity to penetrate the desert pavement and enter the root zone restricts microbial and plant productivity [1, 7]. Consequently, soil organic matter, nitrogen and carbon content are lower in the hyperarid compared to the arid inland region [3, 11, 12]. High soil ionic content from marine aerosol deposition in the coastal region may further limit the bioavailability of fog-derived moisture [12]. While it is clear that fog is a strong driver of lichen and biological soil crust development [8, 13, 14], there has been extensive debate over the role of fog-derived water in supporting the sub-surface soil microbiome [10, 12, 15, 16].

### Microbial trace gas oxidation and alternative survival strategies in dryland environments

It is well-established that the majority of microbial growth and metabolic processes cease at water activities (a_w_) < 0.9, whilst limited function at water activities as low as 0.3 a_w_ occurs in some specialist taxa [17–20]. The Namib Desert is an environment under severe water stress, with average immediate sub-surface (1–2 cm) soil water activities < 0.5 a_w_ [3]. Despite this stress, diverse and metabolically active microbial communities persist within both surface and shallow sub-surface soils [12, 20–23]. It is estimated that over 95% of soil microorganisms exist in slow-growing or dormant states [24, 25], particularly during desiccation stress [19]. Microbial dormancy followed by rapid resuscitation during wetting has been proposed as a key survival strategy in dryland ecosystems including the Namib Desert [22, 26–28].

Microbial trace gas oxidation is increasingly recognised as a generalist strategy to support persistence in both hot and cold desert ecosystems [27, 29–32]. In particular, molecular hydrogen (H_2_) is a ubiquitous and energetically favourable substrate present at ∼530 ppbv throughout the atmosphere, capable of readily diffusing through soil surface layers and cell membranes [27, 30, 33]. In some hyperarid environments, rapid H_2_ uptake and the presence of abundant high-affinity hydrogenases has led to the hypothesis that energy derived from hydrogen oxidation may exceed requirements for cell maintenance, supporting microbial growth via carbon fixation pathways [34–37]. This process, termed atmospheric chemosynthesis, may complement or even surpass photoautotrophic primary production in water-restricted environments with low phototroph abundances. Atmospheric chemosynthesis has been established as a significant contributor to microbial carbon fixation in polar soils [34, 36]. However, microbial trace gas oxidation and its potential contribution to carbon fixation is poorly characterised in hot deserts, particularly in desiccated soils [31, 38].

In this pilot study, the impact of water limitation and regime on microbial atmospheric hydrogen oxidation activity was investigated in desiccated and wetted soils, across a longitudinal aridity gradient in the Namib Desert. The inverse distribution of fog and rainfall-dominated water regimes across the coastal and inland regions of this gradient are proposed to drive divergent microbial community composition and function. Trace gas chemotrophy is predicted to support shallow sub-surface microbial communities in these severely water-limited soils, particularly in the hyperarid coastal zone receiving low-volume fog inputs.

## Materials and Methods

### Soil sampling across the Namib Desert C14 aridity transect

Ten sites were sampled across a 200 km longitudinal aridity transect, referred to henceforth as the C14 transect, spaced at 20 km intervals across a well-defined west-east aridity gradient from the coastal to inland margins of the Namib Desert (Figure 1B). The C14 transect and surrounding soils are divided into three well-defined xeric zones: the “Fog zone” (coastal sites 2–6), “Hyperarid zone” (central sites 8–14) and “Arid zone” (inland sites 16–20). These zones, proposed by Scola et al. (2018) [12], are supported by extensive research on soil physicochemical parameters [11, 12, 16, 39, 40], rainfall, fog and humidity records [2–4, 11, 12, 16, 39], and microbial community analysis [12, 39, 40].

Duplicate (approx. 45 g) shallow sub-surface (1–5 cm) soil samples were aseptically collected into sterile 50 mL tubes from the ten C14 transect sites in April 2023 (Figure 1B, Table 1). Samples were immediately stored at 4 °C after sampling and transported to the Centre for Microbial Ecology and Genomics (CMEG, Pretoria, South Africa), where they were stored at 4 °C until shipment. Samples were imported to the University of New South Wales (NSW, Australia), where half of the bulk soil from each site was stored at −80 °C and the remaining soil stored at 4 °C until use. Soil samples used for hydrogen oxidation and respiratory burst microcosm assays were stored at 4 °C, while DNA extraction and all other analysis was conducted on soils stored at −80 °C.

**Table 1:** Soil moisture metrics and physicochemical properties of C14 transect sampling sites.

| Site | Xeric zone <sup>a</sup> | Lat | Lon | Aridity Index (AI) | Gravimetric SMC% | Annual rainfall (mm) | Water potential (MPa) | Water activity (a<sub>w</sub>) | Sub-surface SMC (m<sup>3</sup>/m<sup>3</sup>) <sup>b</sup> | TC% | TIC% | TOC% | N% | H% | S% <sup>c</sup> | pH |
| --- | --- | --- | --- | --- | --- | --- | --- | --- | --- | --- | --- | --- | --- | --- | --- | --- |
| C14-2 | F | -23.001 | 14.672 | 0.0072 | 1.53 | 16.7 | -100.81 | 0.498 | 0.173 | 0.27 | 0.19 | 0.08 | 0.05 | 0.41 | 2.32 | 6.70 |
| C14-4 | F | -23.018 | 14.860 | 0.0112 | 0.20 | 21.5 | -123.47 | 0.432 | 0.182 | 0.18 | 0.14 | 0.04 | 0.04 | 0.06 | 0.08 | 7.19 |
| C14-6 | F | -23.066 | 15.040 | 0.0196 | 0.82 | 40 | -144.38 | 0.384 | 0.169 | 0.63 | 0.55 | 0.08 | 0.03 | 0.26 | 0.39 | 6.91 |
| C14-8 | HA | -23.143 | 15.209 | 0.0233 | 0.23 | 59 | -162.43 | 0.333 | 0.163 | 0.16 | 0.11 | 0.05 | 0.04 | 0.09 | 0.07 | 7.03 |
| C14-10 | HA | -23.246 | 15.360 | 0.0308 | 0.81 | 62 | -162.02 | 0.346 | 0.163 | 1.03 | 0.89 | 0.14 | 0.05 | 0.30 | 0.01 | 8.05 |
| C14-12 | HA | -23.311 | 15.534 | 0.0426 | 0.42 | 78.8 | -184.07 | 0.312 | 0.162 | 0.38 | 0.18 | 0.20 | 0.07 | 0.26 | nd | 7.87 |
| C14-14 | HA | -23.325 | 15.715 | 0.0456 | 0.49 | 100.5 | -145.68 | 0.384 | 0.163 | 1.05 | 0.83 | 0.22 | 0.05 | 0.27 | nd | 7.53 |
| C14-16 | A | -23.321 | 15.862 | 0.0532 | 0.35 | 133 | -207.39 | 0.269 | 0.166 | 1.13 | 0.87 | 0.25 | 0.08 | 0.24 | nd | 7.71 |
| C14-18 | A | -23.345 | 16.011 | 0.0644 | 0.27 | 169.3 | -231.39 | 0.230 | 0.170 | 0.22 | 0.04 | 0.18 | 0.06 | 0.17 | nd | 8.45 |
| C14-20 | A | -23.245 | 16.143 | 0.0780 | 0.31 | 184.8 | -204.08 | 0.288 | 0.178 | 0.74 | 0.10 | 0.64 | 0.09 | 0.33 | 0.18 | 7.45 |

<sup>a</sup> Xeric zonation is denoted as F = Fog zone, HA = Hyperarid zone, A = Arid zone
<sup>b</sup> Sub-surface (0–10 cm) soil moisture content
<sup>c</sup> nd denotes sulfur content not detected

### Soil moisture analysis

Soil moisture content (SMC%) was calculated using the gravimetric oven-drying method [41]. Triplicate 1.5 g soil subsamples from each site were placed into pre-weighed 20 mL glass vials and oven-dried at 105 °C for 48 h. Sample vials were removed from the oven and cooled to RT in a gas-tight container with silica beads (Ajax Finechem) before re-weighing using an analytical balance (Sartorius AG). SMC% was recorded by calculating the mass of water loss after drying as a percentage of the dry soil weight.

Aridity Index (AI) data for each site was retrieved from the Global Aridity Index and Potential Evapotranspiration Database (Version 3) [6]. Soil water potential and water activity was calculated from previously published datasets that used iButton remote sensing data to record sub-surface soil temperature and relative humidity along the C14 transect [3]. Sub-surface (0–10 cm) soil moisture content was retrieved from the FLDAS Noah Land Surface Model L4 Global Monthly 0.1° × 0.1° dataset [42], subset to a 30-year monthly climatic mean using NASA GES DISC by selecting the SoilMoi00_10cm_tavg variable between 1994–2024. Average annual rainfall data from each site was retrieved from a previously published dataset across the C14 transect [12].

### Soil physicochemical parameters

Total soil carbon (TC), organic carbon (TOC), inorganic carbon (TIC), nitrogen (N), hydrogen (H) and sulfur (S) analysis was performed at the Mark Wainwright Analytical Centre (MWAC, UNSW Sydney, Australia). Duplicate 0.25 g soil samples from each transect site were first passed through a 0.15 mm sieve, and soil particles between 0.15–2 mm were finely ground to < 0.15 mm using a mortar and pestle. Ground soil samples were then oven-dried at 45 °C until constant mass (> 72 h). To quantify TOC, two additional soil samples per site were dried and ground according to the same procedure. Inorganic carbonates were removed by acidification with 1 M HCl, added gradually in excess, and incubated in glass vials at RT for 12 h [43]. Acidified samples were washed with 90 mL sterile deionised water and dried at 45 °C until constant mass (> 120 h). Soil mass loss after acidification was recorded using an analytical balance (Sartorius AG), and TOC was calculated as a percentage of total soil mass. CHNS analysis was performed on all samples via combustion at 1150 °C using a varioMACRO CUBE elemental analyser (Elementar, Germany). TIC was calculated as the difference between TC and TOC content.

Soil pH was measured using 2.5 g soil slurries at a 1:2.5 soil:deionised water ratio, with a pH meter (HI12303, Hanna Instruments). Additional physicochemical parameters at C14 transect sites have been investigated previously [12, 40].

### Microbial community activity metrics

#### Intracellular ATP luminescence assay

The BacTiter-Glo™ Microbial Viability Assay kit (Promega) was used to measure intracellular ATP concentrations in C14 transect soil samples using a modified method optimised for soils [44]. In triplicate, 1 g soil subsamples from each site were added to 4 mL 0.85% NaCl in 15 mL tubes and vortexed at maximum speed for 5 min to produce a soil slurry. Slurry samples were then centrifuged at 180 rcf for 5 min and the supernatant was collected. Triplicate 100 µL volumes of supernatant were used to perform the assay in 96-well white luminescence plates (Greiner), according to manufacturer’s instructions. A standard curve was produced by serial dilution of 100 mM ATP solution (Thermo Fisher Scientific). Background adjusted luminescence values were recorded using a CLARIOstar Plus Microplate reader (BMG Labtech), and calculations were performed in MARS Data Analysis Software v5.02 R3 (BMG Labtech).

#### Respiratory activity assay in dry and wetted soils

Basal respiration in dry soils and respiratory burst after wetting were recorded over 14 days using a custom-built CO_2_ and O_2_ respirometry system (Qubit Systems, Canada). Triplicate 1 g subsamples of C14 transect soils stored at 4 °C were placed into sterile 114 mL glass vials (Glass Vials Australia, NSW) and equilibrated at RT for three days. Following equilibration, soil microcosms were sealed using gas-tight butyl rubber stoppers (Glass Vials Australia, NSW) and incubated in the dark at 30 °C. Heat-killed controls were prepared from each site using 1 g soil autoclaved at 121 °C for 15 min.

Headspace CO_2_ and O_2_ was measured at five timepoints (0, 24, 56, 120, 144 and 168 h) over the one-week basal incubation period. A gas-tight syringe (Trajan Scientific) was used to subsample 1 mL headspace gas from serum vial microcosms. Headspace subsamples were injected through a gas-tight sample port at approximately 100 µL/s into 99.999% N_2_ carrier gas (Coregas, Australia). Using a Q-P103 Gas Pump (Qubit Systems), carrier and sample gases were pumped through a Q-S151 Infrared CO2 analyser and Q-S102 O2 analyser (Qubit Systems) at a constant rate. Flow rate was recorded via a Q-G266 Flow Monitor (Qubit Systems). All readings were recorded using LabQuest Mini sensor data interfaces (Qubit Systems) and headspace CO_2_ and O_2_ concentrations were calculated via the Logger Pro v3.16.2 software.

After one week, each soil microcosm was wetted to approximately 50% water holding capacity, estimated using the filter paper method [45]. Starting headspace CO_2_ concentrations were recorded immediately before wetting, as above, and wetted microcosms were incubated under the same conditions. Repeat measurements were taken after 0.5, 1, 3, 5, 9, 24, 48 and 168 h. The cumulative respiratory burst was calculated for each site and timepoint by normalising against the initial headspace CO_2_ concentration.

### Hydrogen oxidation rates in wet and dry soil microcosms

For dry microcosm H_2_ oxidation assays, five biological replicate soil subsamples (2 g) and a heat-killed control (2 g soil autoclaved at 121 °C for 15 min) were prepared from each site, along with an empty serum vial negative control. Microcosm H_2_ oxidation activity was recorded using gas chromatography as described previously [34, 36]. All soil samples and controls were placed into sterile 114 mL serum vials sealed with gas-tight butyl rubber stoppers and incubated in the dark at 30 °C. Sterile H_2_ gas in synthetic air-balance (BOC, Australia) was injected into the headspace of each serum vial microcosm to achieve a final headspace concentration of 10,000 ppbv. Headspace gas from each microcosm was subsampled in 1 mL volumes at regular intervals using a gas-tight syringe (Trajan Scientific) and H_2_ concentration was measured using a Peak Performer 1 Gas Analyser (Peak Laboratories, USA). For H_2_ oxidation analysis in wet soil microcosms, the same procedure was repeated using five additional 2 g soil subsamples from each site, wetted immediately prior to gas addition using 400 µL sterile ddH_2_O equilibrated to 30 °C.

Sub-atmospheric hydrogen oxidation was confirmed after two consecutive headspace H_2_ readings < 530 ppbv. First order rate constants (*k*) were calculated using linear regression (R^2^ > 0.9). Atmospheric hydrogen oxidation rates were calculated at headspace H_2_ = 530 ppbv with a total microcosm headspace volume of 114 mL at 30 °C and 1 atm. Rates were normalised against total 16S rRNA gene copy numbers per gram soil.

### DNA extraction and quantitative PCR

Genomic DNA was extracted from quadruplicate 0.5 g bulk soil subsamples stored at −80 °C from each site, using the FastDNA™ Spin Kit for Soil (MP Biomedicals), according to manufacturer’s instructions. Extracted DNA was eluted in 70 µL warm (60 °C) UltraPure™ DNase/RNase-free distilled water (Invitrogen) and stored at −80 °C. Quantity and purity of extracted DNA was assessed using both Nanodrop (Thermo Fisher Scientific) and the Qubit™ dsDNA High Sensitivity Assay (Thermo Fisher Scientific). To reduce PCR inhibition, gDNA was further diluted in UltraPure™ DNase/RNase-Free distilled water (Invitrogen) to produce final concentrations between 1.76–20.00 ng/µL.

Extracted gDNA was used for qPCR quantification of total bacterial abundance using the universal 16S rRNA gene degenerate primer set Eub1048f/Eub1194r (Eub1048f; 5’-GTGSTGCAYGGYTGTCGTCA, Eub1194r; 5’ACGTCRTCCMCACCTTCCTC) [46], as described previously [47]. QPCR reaction mixtures were prepared using 10 µL QuantiNova SYBR Green PCR Master Mix (Qiagen, Australia), 0.5 µL of each 40 µM forward and reverse primer (Integrated DNA Technologies), 7 µL UltraPure™ DNase/RNase-free distilled water (Invitrogen), 1 µL of 5 µg/mL T4 Gene 32 (New England Biolabs), and 1 µL diluted template gDNA or positive control. A synthetically designed gene fragment (gBlocks; Integrated DNA Technologies, Australia) containing a representative 16S rRNA gene sequence (MF689012.1) was used to generate a standard curve over five orders of magnitude [47]. All standards, samples and negative template control reactions were performed in triplicate.

Reaction mixtures were added to 96-well thin wall PCR plates (Bio-Rad Laboratories) and sealed using optically clear plate seals (Bio-Rad Laboratories). The thermocycling protocol was completed using a CFX96 Touch™ Real-Time PCR Detection System (Bio-Rad Laboratories). Plates were incubated at 95 °C for 5 min, followed by 38 cycles of 95 °C for 20 s and 60 °C for 50 s. Quantitative fluorescence data was spectrophotometrically collected during the combined extension and annealing step. Following PCR amplification, a melt curve was performed from 50–95 °C, with fluorescence data collected at 0.5 °C steps throughout. Data analysis was conducted using CFX Manager software (Bio-Rad Laboratories). Amplification specificity was confirmed by visual inspection of melt peaks.

### 16S rRNA gene amplicon sequencing of soil microbial communities

Paired-end amplicon sequencing was performed on gDNA samples from all transect sites at the Ramaciotti Centre for Genomics (UNSW, Australia). The barcoded primer pair 515F-Y/806RB (515F-Y; 5’-GTGYCAGCMGCCGCGGTAA, 806RB; 5’-GGACTACNVGGGTWTCTAAT) was used to amplify the V4 region of the 16S rRNA gene using the Illumina MiSeq v2 platform [48, 49]. Raw reads were processed into amplicon sequence variants (ASVs) with DADA2 v1.30.0 [50] in R v4.3.1. Reads with ambiguous bases were discarded, and residual primer sequences were removed with cutadapt v4.3 [51]. Forward and reverse reads were quality filtered using the following parameters: reads matching phiX removed, 2 maximum expected errors allowed, truncation length of 220 and 215 nt, minimum read length of 150 nt, and a read truncation quality threshold of 2. Paired reads were merged with mergePairs() using default parameters and chimeric reads were removed with the removeBimeraDenovo() function, using the consensus method. Taxonomy was assigned to merged reads with the assignTaxonomy() command, implementing the RDP Naive Bayesian Classifier algorithm [52] using SILVA database v138.1 [53]. Processed ASVs assigned to mitochondria or chloroplasts were removed.

ASV sequences were aligned using MAFFT v7.526 [54], and an unrooted phylogenetic tree was built using FastTree v2.1.11 [55] under the GTR+Gamma model [56]. The resulting tree was utilised to calculate weighted UniFrac distance matrices [57] using phyloseq v1.46 [58]. Raw ASV abundance tables were rarefied to the minimum read count using phyloseq v1.46 [58]. Microbial community composition across C14 transect sites was analysed in vegan v2.6-10 [59] using principal coordinates analysis (PCoA) based on weighted UniFrac distances and the rarefied count table. Permutational ANOVA (PERMANOVA) was performed using the adonis2 function in vegan v2.6-10 [59] with 999 permutations, where each explanatory variable was tested individually against the weighted UniFrac dissimilarity matrix.

Differentially abundant taxa were identified using Maaslin2 v1.16.0 [60] from rarefied data aggregated to taxonomic level, excluding unclassified taxa. Linear regression models (analysis method = “LM”) were used with default standardisation settings, and no normalisation was applied. The Fog zone was specified as the reference group for comparisons across xeric zones. The minimum non-zero prevalence threshold was set at 30% and nominal p-values were adjusted via the Benjamini-Hochberg correction method to a significance threshold of qval ≤ 0.20.

### Data analysis and visualisation

All community activity and aridity datasets were analysed in RStudio [61] using R v4.3.1. Data processing was performed using reshape2 v1.4.4 [62], stringr v1.5.1 [63] and readr v2.1.5 [64]. Aridity Index datasets were processed and visualised using the terra v1.8.21 [65] and tidyterra v0.7.0 [66] packages. Statistical analysis was performed using dplyr v1.1.4 [67] and ggpubr v0.6.0 [68]. Correlation analysis was performed using corrplot v0.92 [69].

Mantel correlogram was calculated with vegan’s v2.6-10 [59] mantel.correlog() function between the weighted UniFrac distance matrix and the geographical distances between sampling points. Geographical distances between sampling points were calculated from GPS coordinates using the Haversine method [70] as implemented in the geosphere package v1.5-20 [71]. Optimal number of bins for Mantel correlogram was calculated based on the Freedman-Diaconis rule [72] with the hist.FD() function from the MASS package v7.3-60 [73].

All other data visualisation was performed using ggplot2 v3.5.1 [74], patchwork v1.3.0 [75], pals v1.10 [76], tidyverse v2.0.0 [77] and ggtext v0.1.2 [78].

## Results

### Water availability and soil properties across the C14 transect

The Aridity Index (AI) of sites across the C14 transect increased from west to east, with the lowest recorded at Fog zone site C14-2 (AI = 0.0072), and the highest at Arid zone site C14-20 (AI = 0.0780) (Table 1). Sites C14-2 to C14-14, spanning the Fog and Hyperarid zones, were classed as hyperarid (AI < 0.05), and Arid zone sites C14-16–20 were classed as arid (0.05 ≤ AI < 0.2) [5].

Gravimetric SMC% varied across the transect, ranging from 0.20–1.53% (0.54% ± 0.42%, mean ± SD) (Table 1, Table S1). Average SMC% was highest in the Fog zone (0.85% ± 0.59%) followed by the Hyperarid (0.49% ± 0.25%) and Arid (0.31% ± 0.10%) zones. Average water potential and a_w_ were highest in the Fog zone (water potential = −122.89 ± 17.79 MPa; a_w_ = 0.438 ± 0.047), followed by the Hyperarid (water potential = −163.55 ± 13.64 MPa; a_w_ = 0.344 ± 0.026) and Arid (water potential = −214.28 ± 12.17 MPa; a_w_ = 0.262 ± 0.024) zones. AI and rainfall were both negatively correlated with water potential (*p* ≤ 0.001) and a_w_ (*p* ≤ 0.01) (Figure S1). Gravimetric SMC% was positively correlated with a_w_ (ρ = 0.67, *p* ≤ 0.05). Sub-surface (0-10 cm) soil moisture content was not significantly correlated with any other moisture metric (*p* > 0.05).

Soil pH ranged from slightly acidic to alkaline (6.70–8.45) and was highest in the Arid zone (7.87 ± 0.42, mean ± SD), followed by the Hyperarid (7.62 ± 0.39) and Fog (6.93 ± 0.20) zones (Table 1). Total carbon (TC) content was low (0.16–1.13%) across the transect, and inorganic carbon (TIC) represented the majority of soil carbon in all sites except C14-12, 18 and 20. Total nitrogen (N) and organic carbon (TOC) content were lowest in the Fog zone (N = 0.04% ± 0.01%, TOC = 0.06% ± 0.02%) and increased in the Hyperarid (N = 0.05% ± 0.01%, TOC = 0.15% ± 0.07%) and Arid (N = 0.07% ± 0.01%, TOC = 0.36% ± 0.20%) zones. TOC, N and pH were significantly (*p* ≤ 0.05) lower in the Fog zone compared to the rest of the transect and were significantly positively correlated with AI (Figure S1).

### Microbial activity and abundance biomarkers across xeric zones

Microbial activity and 16S rRNA gene abundances increased from west to east across the Fog, Hyperarid and Arid zones of the C14 transect (Figure 2). Total 16S rRNA gene copy numbers ranged from 3.82 × 10^5^ g^-1^ soil at Fog site C14-2 to 3.10 × 10^7^ g^-1^ soil at Arid site C14-16 (Figure 2A, Table S2). Average 16S rRNA gene copy number increased significantly (*p* ≤ 0.01) between each xeric zone as AI and rainfall increased (Figure S2A). The lowest average 16S rRNA gene abundance was observed in the Fog zone (1.04 × 10^6^ copies g^-1^), increasing 9-fold in the Hyperarid zone (9.15 × 10^6^ copies g^-1^) and by an order of magnitude in the Arid zone (2.36 × 10^7^ copies g^-1^).

**Figure 2:**
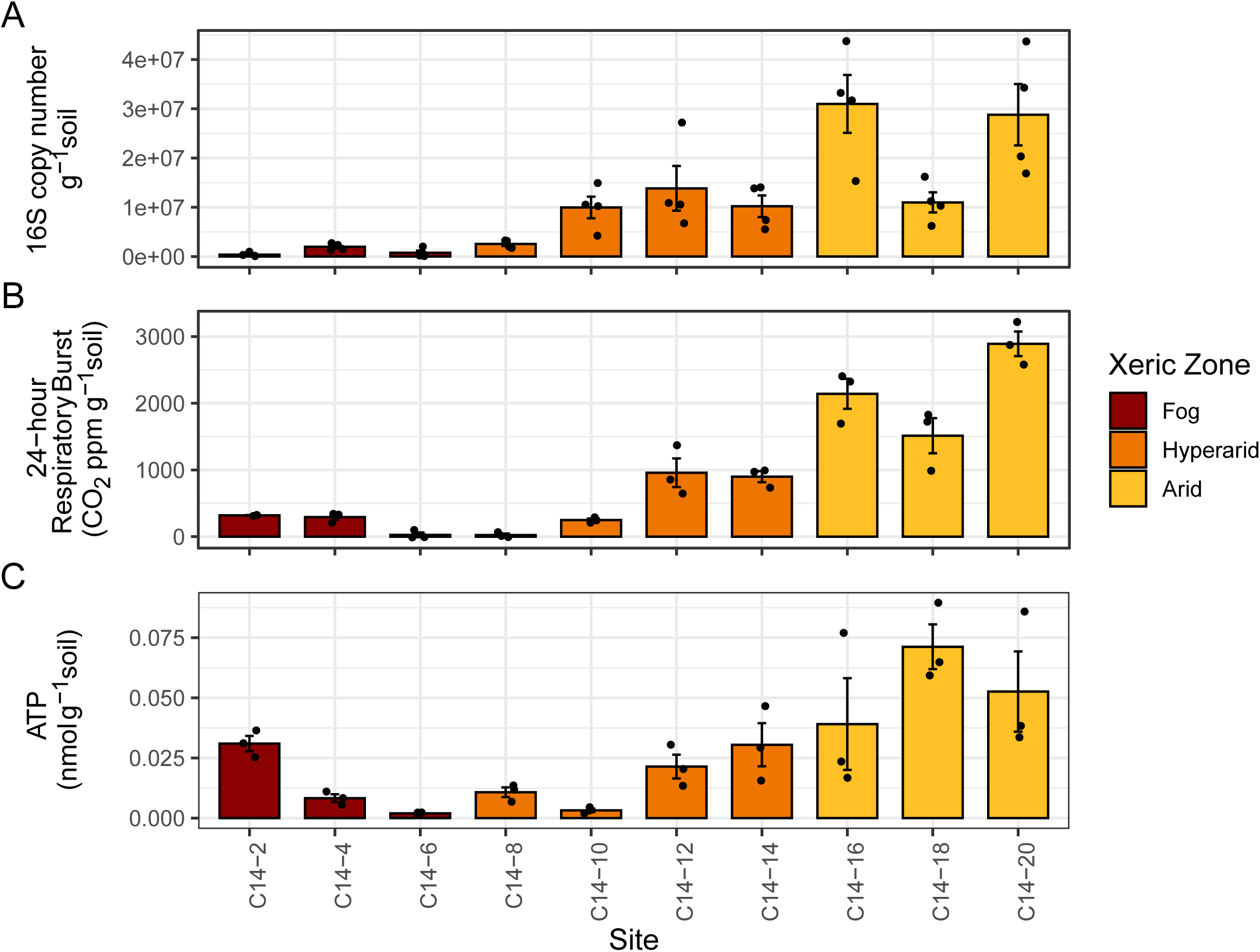
Soil microbial activity and abundances across the Namib Desert C14 transect. Data points represent biological replicates (n=4 for A, n=3 for B and C) and error bars represent standard error. A) Total 16S rRNA gene copy numbers per site. Total copy number increased significantly (p < 0.01) between each xeric zone and was lowest in the Fog zone. B) Cumulative CO2 respiratory burst 24 h after moisture addition per site. Cumulative respiratory burst increased significantly (p < 0.05) between each xeric zone. B) Intracellular ATP concentrations per site. Average ATP was not significantly (p > 0.05) different between the Fog and Hyperarid zone.

Similarly, cumulative 24-hour respiratory burst values ranged from 23.6 ppmv CO_2_ g^-1^ soil (Hyperarid site C14-8) to 2891.0 ppmv CO_2_ g^-1^ soil (Arid site C14-20) (Figure 2B, Table S3-4) and increased significantly (*p* ≤ 0.05) between each xeric zone (Figure S2B). The average 24-hour cumulative respiratory burst across the Fog, Hyperarid and Arid zones was 213.2, 532.1 and 2181.9 ppmv CO_2_ g^-1^ soil, respectively. Intracellular ATP concentrations ranged from 0.002 nmol g^-1^ soil at Fog site C14-6 to 0.071 nmol g^-1^ soil at Arid site C14-18 (Figure 2C, Table S5). Whilst Arid zone sites recorded significantly (*p* ≤ 0.01) higher ATP concentrations than Fog and Hyperarid zone sites, average ATP concentrations between the Hyperarid and Fog zones were not significantly different (*p* > 0.05) (Figure S2C).

Within the Fog zone, average ATP concentrations in sites C14-2 and C14-4 were 15- and 4-fold higher, respectively, compared to site C14-6. Aridity Index was significantly positively correlated with total 16S rRNA gene copy numbers (ρ = 0.83, *p* ≤ 0.01), intracellular ATP (ρ = 0.76, *p* ≤ 0.05) and 24 h respiratory burst (ρ = 0.90, *p* ≤ 0.001) (Figure S1). Total 16S copy numbers were significantly positively correlated with cumulative 24 h respiratory burst (ρ = 0.92, *p* ≤ 0.001) but not intracellular ATP content (*p* > 0.05).

### Trace gas oxidation rates in Namib Desert soil microcosms

In dry soil microcosms, two Fog zone sites (C14-2 and C14-4) exhibited rapid H_2_ oxidation activity to sub-atmospheric (< 530 ppbv) levels (Figure 3A). The remaining dry microcosms demonstrated slow H_2_ oxidation that did not reach sub-atmospheric levels after 1,500 h (∼2 months) (Figure S3A). In contrast, soil wetting resulted in immediate (< 30 min) and rapid H_2_ oxidation activity across all ten transect sites (Figure S3B), with sub-atmospheric hydrogen oxidation activity observed in all moisture-stimulated microcosms (Figure 3B).

**Figure 3:**
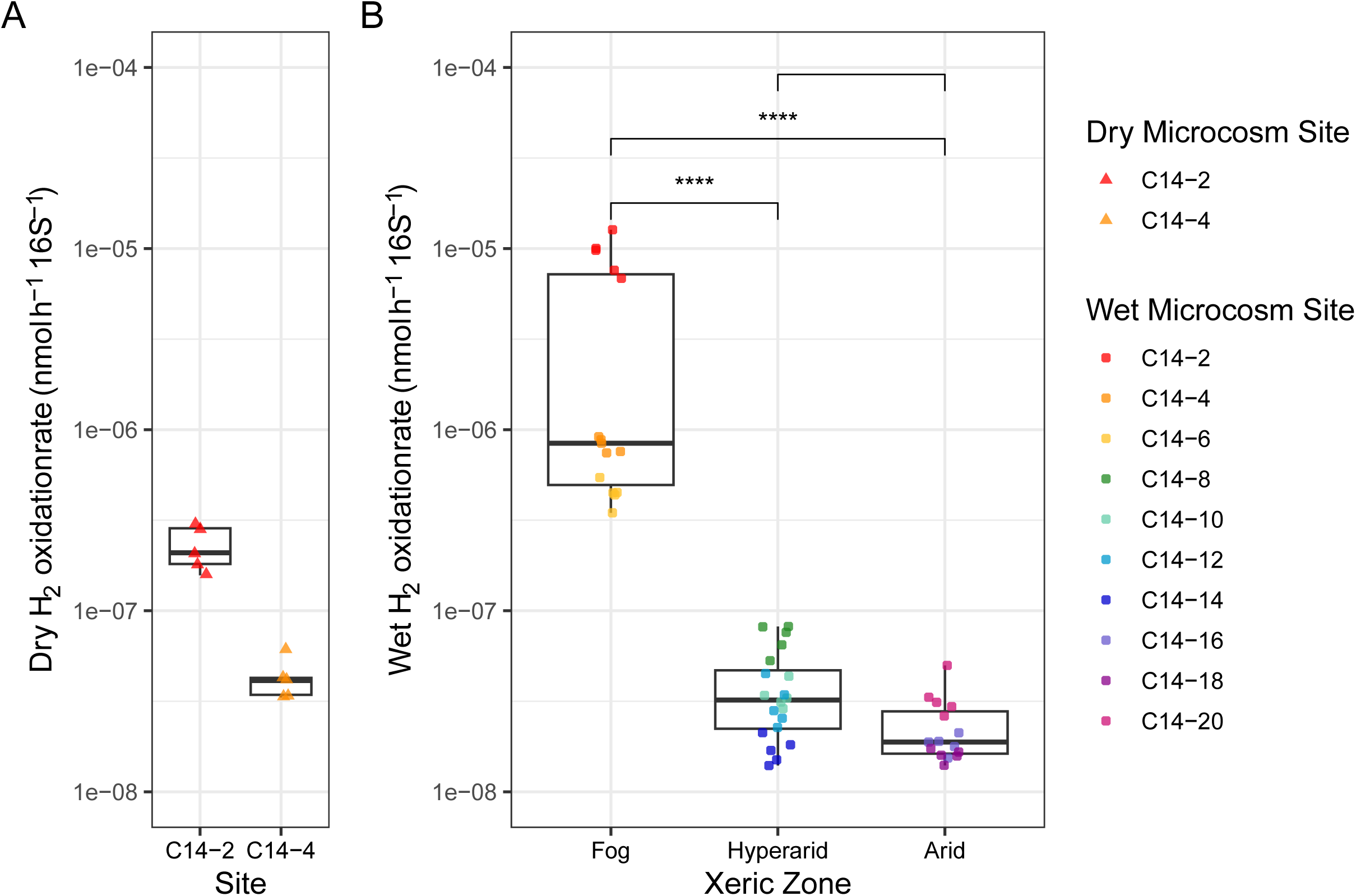
Atmospheric hydrogen oxidation rates in dry and wetted soil microcosms. Rates were calculated at atmospheric hydrogen concentration = 530 ppbv and normalised to average 16S rRNA gene copy numbers per site. Significance thresholds were assigned as: p ≤ 0.05 = *, p ≤ 0.01 = **, p ≤ 0.001 = ***, p ≤ 0.0001 = ****. Data points shown represent n=5 biological replicates. A) Atmospheric H2 oxidation rates in dry soil microcosms from two Fog zone sites. All remaining dry soil microcosms did not oxidise hydrogen to below 530 ppbv after 1,500 hours. B) Atmospheric H2 oxidation rates in wetted microcosms.

At atmospheric headspace H_2_ concentrations, dry soil microcosms from Fog sites C14-2 and C14-4 oxidised hydrogen at average rates of 2.27 × 10^-7^ and 4.27 × 10^-8^ nmol h^-1^ 16S copy^-1^, respectively (Table S6). Both C14-2 and C14-4 microcosms carried out significantly (*p* ≤ 0.0001) accelerated atmospheric H_2_ oxidation rates following moisture stimulation, increasing over 30-fold compared to dry microcosm rates (Figure S4). In contrast, all wet soil microcosms demonstrated rapid atmospheric H_2_ oxidation rates, ranging from 1.59 × 10^-8^ nmol h^-1^ 16S copy^-1^ at Arid site C14-18 to 9.39 × 10^-6^ nmol h^-1^ 16S copy^-1^ at Fog site C14-2 (Figure 3B, Table S7). In moisture-stimulated microcosms, the fastest H_2_ oxidation rates were observed in Fog zone soils (average = 3.55 × 10^-6^ nmol h^-1^ 16S copy^-1^), while atmospheric H_2_ oxidation rates decreased significantly (*p* ≤ 0.0001) in the Hyperarid (3.84 × 10^-8^ nmol h^-1^ 16S copy^-1^) and Arid (2.28 × 10^-8^ nmol h^-1^ 16S copy^-1^) zones (Figure 3B).

### Microbial community composition along the Namib Desert C14 transect

Soils were dominated by the bacterial phyla *Actinobacteriota*, *Proteobacteria*, *Chloroflexi* and *Bacteroidota*, and the archaeal phylum *Crenarchaeota*, together accounting for > 77.3% of the microbial community (Figure 4A, Table S8). The relative abundance of *Actinobacteriota* decreased significantly (*p* ≤ 0.001) across the transect as AI increased (Figure S6), decreasing from 42.3% in the Fog zone to 22.1% in the Arid zone (Figure 4A). Conversely, the relative abundance of *Proteobacteria* increased significantly (*p* ≤ 0.005) from 11.5% in the Fog zone to 26.7% as the most abundant phylum in the Arid zone. The five most abundant families were *Rubrobacteriaceae* (13.4%, *Actinobacteriota*), *Nitrososphaeraceae* (8.3%, *Crenarchaeota*), *Beijerinckiaceae* (7.7%, *Proteobacteria*), *Chloroflexi* family AKIW781 (4.9%) and *Geodermatophilaceae* (3.9%, *Actinobacteriota*) (Figure 4B).

**Figure 4:**
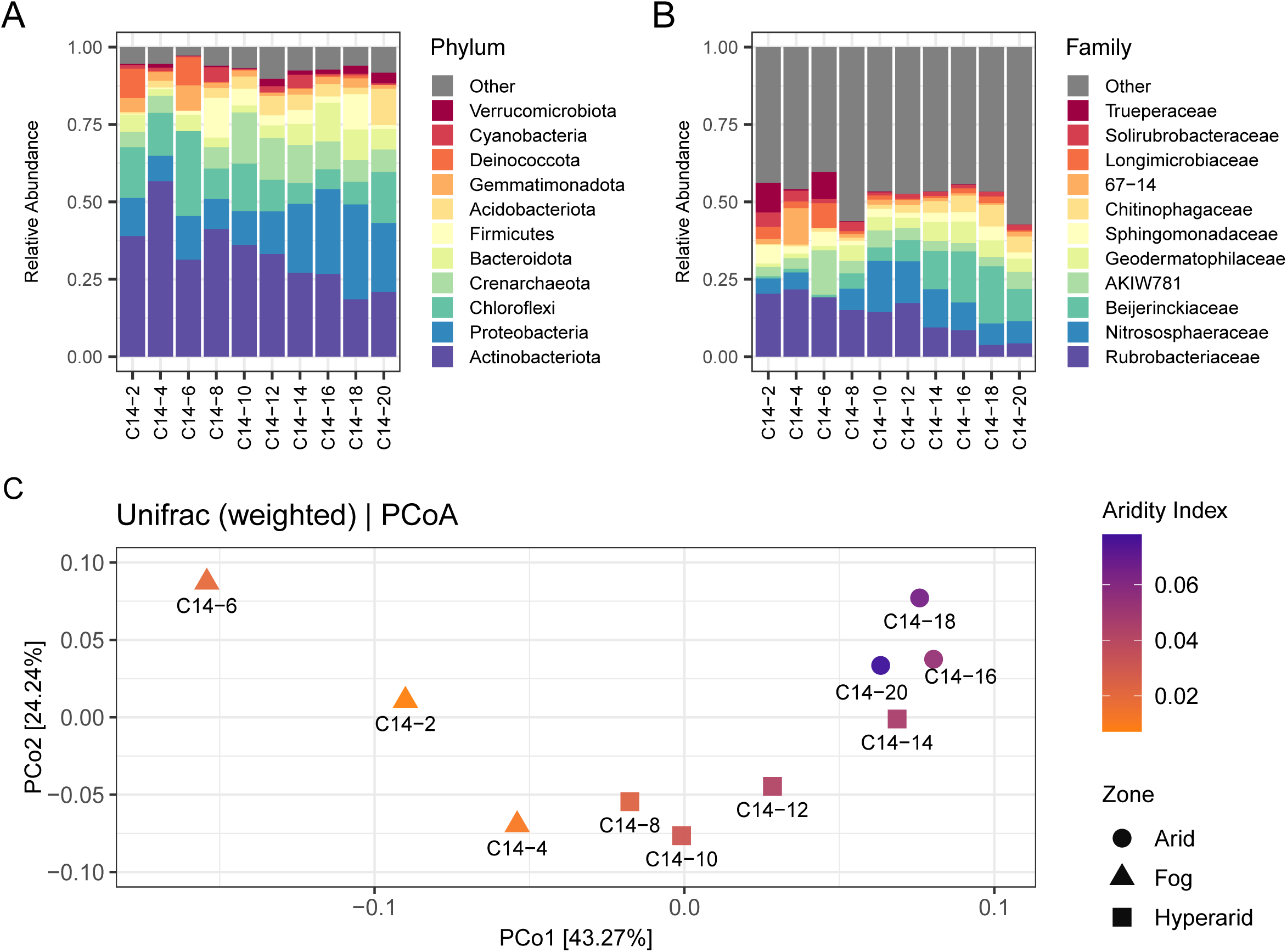
Bacterial and archaeal community composition and drivers across the Namib Desert C14 transect. A) Relative community composition per site aggregated to phylum level. B) Relative community composition per site aggregated to family level. C) Principal coordinates analysis (PCoA) of microbial community composition based on weighted UniFrac distances. PCo1 and 2 explain 43.27% and 24.24% of total variance, respectively. Xeric zone is indicated by shape and transect sites are coloured according to Aridity Index (Table 1).

PCoA ordination analysis explained 67.51% of total variance across the first two axes and showed clear seperation of microbial communities by xeric zone across the first axis (Figure 4C). Arid sites C14-16–20 clustered tightly, whereas Hyperarid sites C14-8–14 showed greater dispersal. Fog zone sites C14-2–6 exhibited the highest dispersion across both axes, indicating relatively higher compositional variability compared to the other zones. AI was the most significant (*p* ≤ 0.001) explanatory variable, explaining 34.78% of variance in community structure (Table S9). Community composition was also significantly associated with rainfall (*p* ≤ 0.005, R^2^ = 0.35), nitrogen (*p* ≤ 0.01, R^2^ = 0.29), water potential (*p* ≤ 0.01, R^2^ = 0.29), pH (*p* ≤ 0.05, R^2^ = 0.25) and water activity (*p* ≤ 0.05, R^2^ = 0.27). The Mantel correlogram showed significant positive spatial autocorrelation (r = 0.37, *p* < 0.01) between sites up to approximately 20 km apart (i.e. immediately consecutive sites) (Figure S5).

Between 20-60 km, correlations were weak (r < 0.2) and non-significant (*p* > 0.05), indicating that geographical distance alone does not explain community differences. At distances of ∼110 km, sites were more dissimilar than expected by chance (r = −0.33, p < 0.05), while non-significant (*p* > 0.5) negative correlations were observed at all other distances.

In total, 129 taxa (13 phyla, 4 classes, 26 orders, 37 families and 49 genera) were significantly (qval ≤ 0.20) differentially abundant across xeric zones (Figure 5, Figure S7, Table S10). At the phylum level, the relative abundance of *Acidobacteriota*, *Abditibacteriota, Firmicutes*, *Proteobacteria, Bacteroidota* and *Crenarchaeota* increased significantly (qval ≤ 0.20) in the Arid compared to the Fog zone. Similarly, seven phyla were enriched in the Hyperarid zone relative to the Fog zone: *Entotheonellaeota, Firmicutes*, *Nitrospirota*, *Acidobacteriota*, *Abditibacteriota*, *Armatimonadota* and *Crenarchaeota*. Conversely, the Fog zone contained significantly (qval ≤ 0.20) greater relative abundances of *Gemmatimonadota*, *Actinobacteriota*, *Deinococcota* and *Bdellovibrionota*.

**Figure 5:**
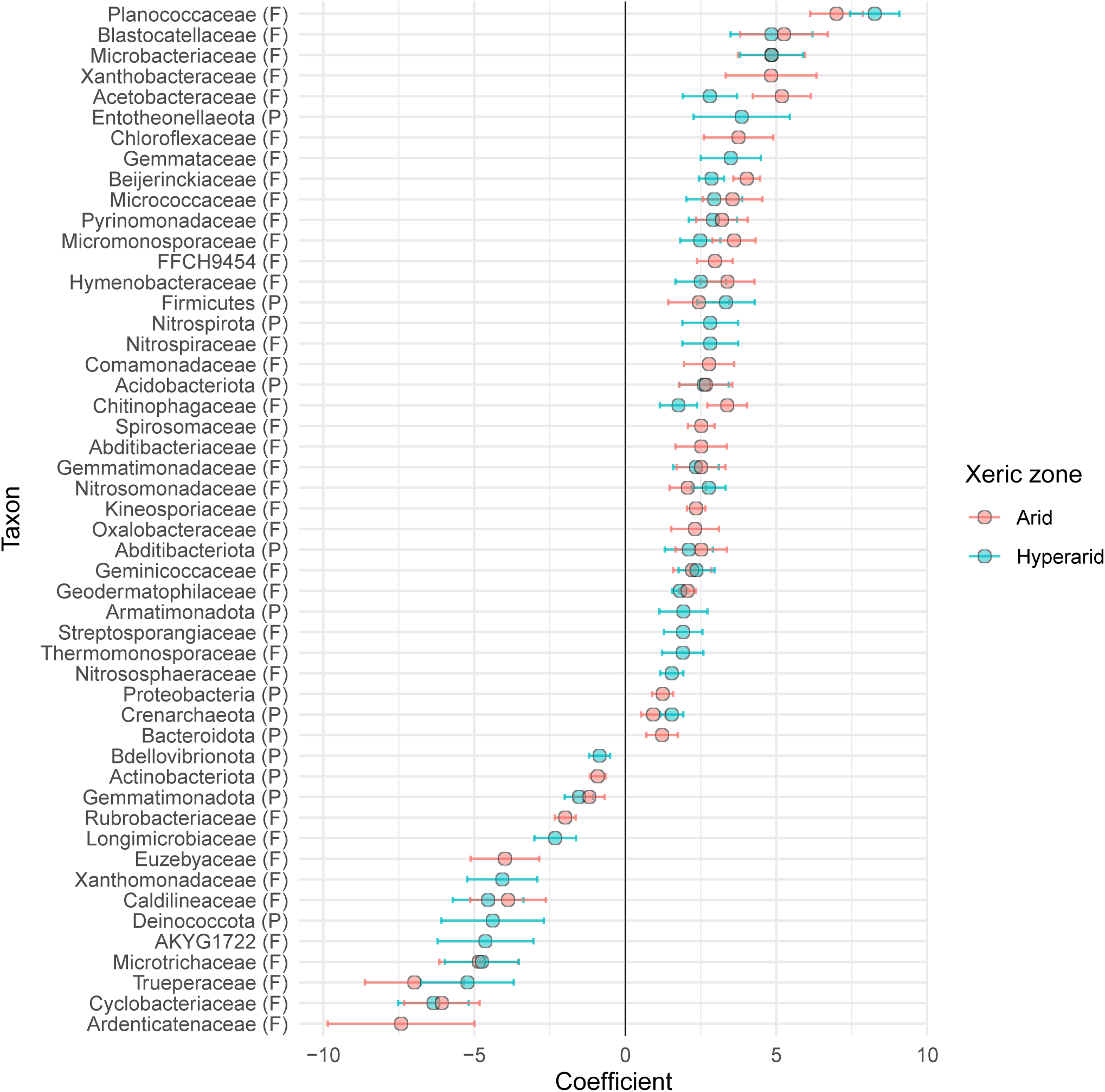
Differentially abundant taxa across xeric zones of a Namib Desert aridity transect, relative to the Fog zone. Coefficient is a measure of the difference in relative abundance of taxa in Hyperarid and Arid zone sites, relative to Fog zone sites. Negative values indicate greater relative abundance within the Fog zone. Only coefficients with adjusted q-values ≤ 0.20 are shown. Error bars represent standard error. Taxonomic levels are represented as follows: (P) = phylum, (F) = family.

Eight of the 12 families with total relative abundance ≥ 1.5% were significantly (qval ≤ 0.20) differentially abundant between xeric zones (Figure 5). Of these, relative abundances of *Chitinophagaceae, Geodermatophilaceae, Beijerinckiaceae* and *Pyrinomonadaceae* were significantly (qval ≤ 0.20) greater within both the Arid and Hyperarid zones relative to the Fog zone. In contrast, *Nitrososphaeraceae* were significantly more abundant in the Hyperarid but not the Arid zone. Within the Fog zone, *Trueperaceae* were present at a greater relative abundance compared to both other zones, while *Longimicrobiaceae* and *Rubrobacteriaceae* had a greater relative abundance compared to the Hyperarid or Arid zone, respectively.

Across the transect, > 43.5% of ASVs remained unclassified at the genus level, with over half of total ASVs unclassified in sites C14-4, C14-6 and C14-20 (Table S8). Seven classified genera were significantly (qval ≤ 0.20) enriched in the Fog zone (Figure S7, Table S10). The relative abundance of *Truepera, Oxalicibacterium* and *Amaricoccus* was greater in the Fog zone relative to both other zones, while *Conexibacter,* Ellin6055 and *Lysobacter* were enriched compared to the Hyperarid zone only, and *Rubrobacter* relative to the Arid zone. An additional five genera were strongly differentially abundant (coef > 5.0), all enriched within the Arid zone: *Flaviaesturariibacter, Planococcus, Edaphobaculum, Blastocatella* and *Massilia*.

## Discussion

Differences in moisture regime and aridity across xeric zones of the Namib Desert are well-established drivers of microbial community composition and function [12, 39, 40]. In this pilot study, high-affinity H_2_ oxidation and rapid resuscitation following soil wetting were identified as key mechanisms likely to be supporting sub-surface soil microbial communities across this aridity gradient. Dry soil microcosms from the hyperarid Fog zone exhibited rapid hydrogen oxidation to sub-atmospheric concentrations without moisture stimulation (Figure 3A). In low-abundance Fog zone sites capable of H_2_ oxidation in dry conditions, elevated intracellular ATP concentrations compared to Fog zone site C14-6 were also observed, providing evidence of a continuously active and strongly desiccation-resistant microbial community.

### Trace gas oxidation supports microbial communities in hyperarid Namib Desert soils

Across the Namib Desert C14 transect, AI and average annual precipitation were highest in the Arid zone (Table 1), with significantly (*p* ≤ 0.005) higher total microbial abundances, intracellular ATP and respiratory burst observed compared to the Hyperarid and Fog zones (Figure 2, Figure S2). Despite this reduced biomass, dry soils from Fog zone sites C14-2 and C14-4 demonstrated a remarkable capacity for rapid, high-affinity (530 ppbv) hydrogen oxidation without water addition, at rates of 2.27 × 10^-7^ and 4.27 × 10^-8^ nmol h^-1^ 16S copy^-1^, respectively (Figure 3A). These rates exceed biomass-normalised H_2_ oxidation rates recorded in hyperarid Negev Desert soils by over two orders of magnitude [38], and contrast with previous findings from hot desert soil studies where sub-atmospheric H_2_ oxidation activity was strongly limited in the absence of soil wetting [29, 31]. Although even faster atmospheric H_2_ oxidation rates have been recorded in Antarctic desert soils [34, 36], oxidation rates in these Fog zone sites were comparable to those in cold, arid environments such as the high Arctic [36] and Andean Altiplano [79], as well low-temperature Antarctic microcosms [35].

In these areas, energy derived from atmospheric hydrogen oxidation is predicted to sustain microbial community biomass and potentially support hydrogenotrophic growth. Given the extremely limited organic carbon availability (0.04–0.08%) and infrequent rainfall within the Fog zone (Table 1) [3, 12], trace H_2_ oxidation is likely to be a key contributor to cellular energy requirements.

Higher soil moisture content and water potential within the Fog zone due to regular fog inputs and humid coastal airflow [3] may have contributed to this unusual capacity for high-affinity hydrogen oxidation in dry soils. However, these soils were severely water-restricted, with average annual water availability < 0.5 a_w_ [3] and gravimetric SMC% as low as 0.20% at site C14-4 (Table 1). Prior transcriptomic investigations on Namib Desert Fog zone soils have revealed active transcription of genes associated with major metabolic pathways including carbon and nitrogen assimilation, resuscitation and replication, even at exceptionally low water activities (< 0.28 a_w_) [20]. Furthermore, both Fog zone sites capable of sub-atmospheric H_2_ oxidation under dry conditions also exhibited elevated intracellular ATP concentrations (Figure 2C), a known byproduct of hydrogen oxidation [80], compared to Fog site C14-6. These findings support the growing body of evidence that some microbial cells in hot desert soils retain significant metabolic functionality, including trace gas oxidation, during severe desiccation [19].

### Soil wetting results in rapid microbial resuscitation and hydrogen oxidation

Rapid resuscitation following re-wetting is a dominant microbial survival strategy in global dryland soils, with the majority of microbial growth restricted to short windows during intermittent rain events [3, 22, 26, 28, 81]. In this study, moisture stimulation resulted in immediate (< 30 min) H_2_ uptake in soil microcosms from all Namib Desert C14 transect sites (Figure S3B). In similar desiccated dryland samples, soil wetting has prompted rapid (< 15 min) resuscitation and upregulation of metabolic activity, including H_2_ oxidation [26, 28, 31]. Consistent with observations of Australian and Negev Desert microcosms [26, 31, 38], soil wetting significantly (*p* < 0.0001) increased hydrogen oxidation rates over 30-fold (Figure S4) and resulted in sub-atmospheric H_2_ oxidation activity in all microcosms (Figure 3B). In wetted soil microcosms, biomass-adjusted H_2_ oxidation rates were highest in Fog zone sites and decreased significantly (*p* < 0.01) across xeric zones as aridity decreased (Figure 3B).

These findings support the hypothesis that trace gas oxidation is a survival strategy associated with aridity and organic carbon limitation, complementing phototrophic and heterotrophic growth in moisture-limited environments [38, 82]. Many *Actinobacteriota* regulate dormancy and resuscitation of diverse bacterial lineages within the soil microbiome through the production of extracellular resuscitation promoting factors (RPFs) [83, 84], and are also implicated in trace gas oxidation [32]. The high relative abundance of *Actinobacteriota* taxa across the transect (18.5–56.7%), particularly within the Fog zone (Figure 4A), may have contributed to this rapid increase in H_2_ oxidation activity after wetting.

### Impact of water regime and availability on Namib Desert microbial community **composition**

Water availability is a major driver of microbial abundance, diversity, and activity in terrestrial environments, with the diversity and abundance of soil bacteria and fungi decreasing as aridity increases [15, 19, 85, 86]. Photosynthetic growth is strongly dependent on bioavailable water at minimum thresholds of 0.8–0.9 a_w_, with *Cyanobacteria* particularly inhibited by desiccation [9, 17]. Although *Cyanobacteria* have been identified as key taxa in the hypolithic communities that colonise the underside of quartz rocks within the Namib Desert, the relative abundances of these taxa are markedly reduced in open soils and biocrusts [9, 23, 87]. This trend was reflected across the C14 transect, where *Cyanobacteria* comprised 0.2–4.7% of the total community (Figure 4A). Instead, sites were dominated by *Actinobacteriota* (18.5–56.7%), *Proteobacteria* (8.2–30.6%) and *Chloroflexi* (6.5–27.5%) lineages (Figure 4A). These metabolically flexible phyla have been linked with trace gas oxidation and the genetic capacity for atmospheric chemosynthesis, and are proposed to support primary production in water-limited environments [34–36, 38, 47].

Across the transect, the ratio between *Actinobacteriota* and *Proteobacteria* varied by xeric zone, with *Actinobacteriota* significantly (qval ≤ 0.20) decreasing in relative abundance as AI increased (Figure S6). In the Fog zone, *Actinobacteriota* were dominant (42.3% relative abundance; Figure 4A), with *Rubrobacteriaceae* highly abundant across the transect (3.7–21.7%, Figure 4B) and significantly enriched in the Fog zone at both the family and genus (*Rubrobacter*) level (Figure 5, Figure S7, Table S10). *Rubrobacteriaceae* are associated with strong resistance against UV radiation, desiccation, heat and oxidative stress, as well as extensive DNA repair potential and mixotrophic growth strategies [26, 81, 88]. Furthermore, *Rubrobacteriaceae* are key active taxa in global dryland soil microbial communities, including in the Namib [20, 28], Negev [26, 81] and Atacama Deserts [86, 89]. High-affinity oxygen-tolerant [NiFe]-hydrogenases have been identified in the genomes of dryland-associated *Rubrobacteriaceae* [26, 36, 38, 81], and hydrogenase mRNA transcripts assigned to *Rubrobacteriaceae* MAGs have been recovered from microcosms capable of trace H_2_ oxidation under wet and dry conditions [26, 31]. In polar and hot deserts, *Rubrobacteriaceae* are associated with mixotrophic growth due to the genetic capacity to use energy derived from trace hydrogen oxidation and rhodopsin-based light harvesting to support persistence or growth during starvation [35, 81]. Several other differentially abundant Fog zone taxa present at lower relative abundances are also associated with hydrogen chemotrophy, including *Conexibacter* (*Actinobacteria*) [36, 90], *Euzebyaceae* (*Actinobacteria*) [36, 91], *Xanthomonadaceae* (*Proteobacteria*) [35, 36] and *Trueperaceae* (*Deinococcota*) [36]. These hydrogenotrophic and desiccation resistant taxa, particularly the highly abundant *Rubrobacteriaceae,* likely contribute to the significantly faster rates of H_2_ oxidation observed within the Fog zone.

In the Arid zone, the differentially abundant families *Beijerinckiaceae* (*Proteobacteria*) and *Chitinophagaceae* (*Bacteroidota*) constituted > 20.9% of the microbial community (Figure 5, Figure 4B). Within the dominant *Beijerinckiaceae* family, > 98.5% of classified ASVs were assigned to two genera: *Microvirga* and *Psychroglaciecola* (Table S8). These heterotrophic nitrogen-cycling genera are associated with bryophytes and rhizosphere root nodules across a variety of arid environments, including the Namib Desert [21, 92–96]. Many *Microvirga* species have the capacity to utilise plant-derived substrates for growth and synthesise plant-growth promoting compounds [97–99] and have been identified as highly abundant in the rhizosphere of Namib Desert plants such as *Sesuvium sesuvioides* and *Stipograstis* species [21, 93]. Heterotrophic growth strategies may be more prevalent in the Arid zone compared to fog-dominated areas due to the capacity of rainfall inputs to saturate soil and facilitate dispersion of organic matter, as well as the higher TOC (0.18–0.64%) and N (0.06–0.09%) content in the Arid zone (Table 1) [15]. The production of plant-growth promoting compounds has also been reported in the differentially enriched Arid zone *Chitinophagaceae* family, which are abundant within the rhizosphere of Namib Desert *Acanthosicyos horridus* shrubs and are associated with degradation of plant-derived cellulose and chitin [100–102]. These microorganisms may contribute to supporting plant growth in the nutrient and moisture limited Arid region of the Namib Desert.

Across the transect, a high proportion of ASVs belonged to unclassified clades, with > 26.9% unclassified at the family level and > 43.6% at the genus level (Table S8). This high relative abundance of “microbial dark matter” has been observed across global drylands [103], representing a significant and underexplored microbial resource. Further research into these uncharacterised taxa is recommended to support a greater understanding of the complex microbially mediated processes within desert environments.

### Microbial hydro-genesis in Namib Desert sub-surface soils

Hydrogen oxidation, in addition to supporting energy requirements for basal cellular metabolism [27, 33, 104], has been proposed to contribute to intracellular water production through a process termed microbial hydro-genesis [15, 19, 35]. In this study, the determination of substantial H_2_ oxidation rates in dry soil microcosms, normalised against cell numbers (estimated as 16S gene copies), allows for preliminary quantitative assessment of intracellular water input derived from trace hydrogen oxidation.

Using the fastest H_2_ oxidation rate determined in unwetted soil microcosm experiments (2.27 × 10^-7^ nmol h^-1^ 16S copy^-1^, soil sample C14-2) (Figure 3A) and making the assumption that cells contain single genomes and a single 16S gene copy per genome, we estimate H_2_ oxidation (equimolar with H_2_O production) to be 2.27 × 10^-7^ nmol h^-1^ cell^-1^, equivalent to 4.09 fg H_2_O produced h^-1^ cell^-1^. Given a typical saturated cellular water content of 516 fg (for *E. coli*, [105]), this contributes approximately 0.79% of total (saturated) cell water per hour. In desiccated cells, such as those from site C14-4 (SMC% = 0.20%, a_w_ = 0.43; Table 1), this contribution would be proportionally higher. While we acknowledge the multiple assumptions built into this calculation, particularly those related to 16S gene copies and cellular water contents, we argue that this calculated value of hydro-genesis derived from atmospheric hydrogen oxidation in unwetted, low-moisture soils is significant, providing support to the suggestion that this process contributes to cellular water budgets. This hydrogenic contribution, while unlikely to be functionally significant in water replete cells, may be vitally important to cellular function in moisture-limited environments [15, 19, 37].

### Conclusions

In this study, the critical role of trace gas oxidation, particularly high-affinity hydrogen oxidation, in sustaining microbial communities in hyperarid Namib Desert soils is highlighted. Rapid sub-atmospheric H_2_ oxidation activity in unwetted moisture-limited soils is proposed to contribute to cellular ATP and water budgets, supporting a continuously active desiccation resistant microbial community. In hyperarid, fog-dominated soils, taxa implicated in trace gas oxidation dominated, while heterotrophic taxa were differentially abundant within the Arid zone. Emerging evidence of interconnected phototrophic, heterotrophic and hydrogenotrophic growth strategies in hot and polar desert soils supports the need for broader studies that link water regime to functional capacity and primary production dynamics in these complex ecosystems.

## Acknowledgements

This research was supported by an Australian Government Research Training Program (RTP) Scholarship awarded to DZT, the Australian Research Council (ARC) Discovery Project (DP240102658) awarded to BCF and a National Research Foundation of South Africa grant (137954) awarded to DAC. We thank the Gobabeb-Namib Research Institute (http://gobabeb.org/) in Namibia for their hospitality and field assistance. Field research was conducted under a research permit RPIV00112018, issued by the National Commission on Research, Science and Technology, and supported by the Ministry of Environment, Forestry and Tourism in Namibia. This research includes computations using the computational cluster Katana supported by Research Technology Services at UNSW Sydney (DOI: 10.26190/669X-A286).

## CRediT author contribution statement

**DZT** (Formal analysis, Investigation, Methodology, Visualisation, Writing – original draft, Writing – review & editing), **PHL** (Conceptualisation, Writing – review & editing), **XVC** (Formal analysis, Data curation, Visualisation, Writing – review & editing), **AER** (Methodology, Writing – review & editing), **TL** (Formal analysis, Methodology), **NMDL** (Formal analysis, Writing – review & editing), **GMK** (Project administration, Resources), **DAC** (Conceptualisation, Funding acquisition, Project administration, Resources, Supervision, Writing – review & editing), **BCF** (Conceptualisation, Funding acquisition, Project administration, Resources, Supervision, Writing – review & editing). All authors contributed to the final manuscript.

## Data availability

Raw amplicon sequencing data are deposited upon publication in ENA under BioProject accession PRJEB97070. All remaining data generated or analysed during this study are included in this published article and its supplementary information files.

