## Supplementary Figures for "Trace gas oxidation supports sub-surface microbial communities across Namib Desert fog and aridity gradients"

***for***

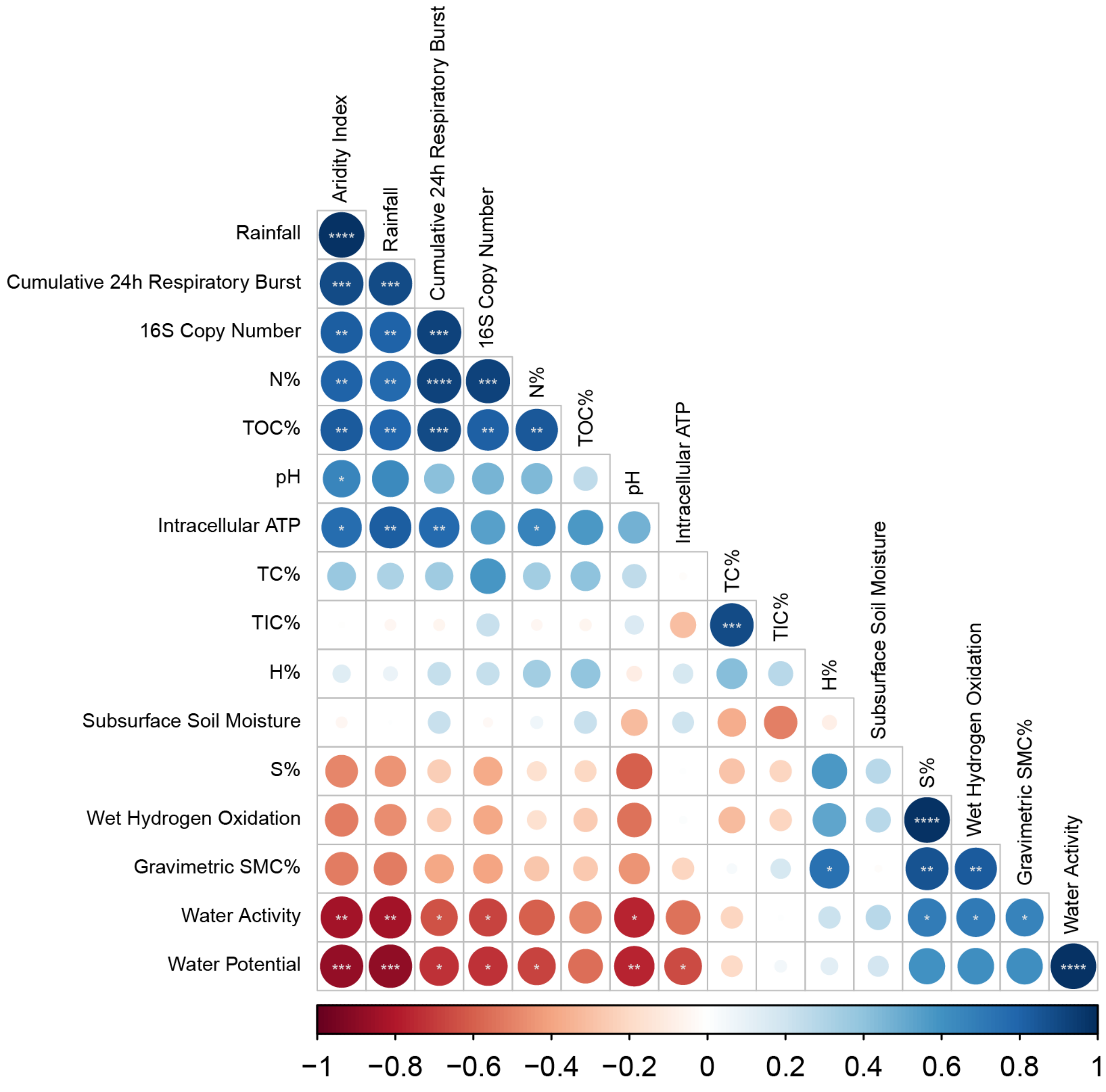


***Figure S1: Correlation matrix of soil moisture metrics and biomarkers of microbial community activity and abundance**. Significance levels are assigned as follows: $p \leq 0.05$ = *, $p \leq 0.01$ = **, $p \leq 0.001$ = ***, $p \leq 0.0001$ = ****.*

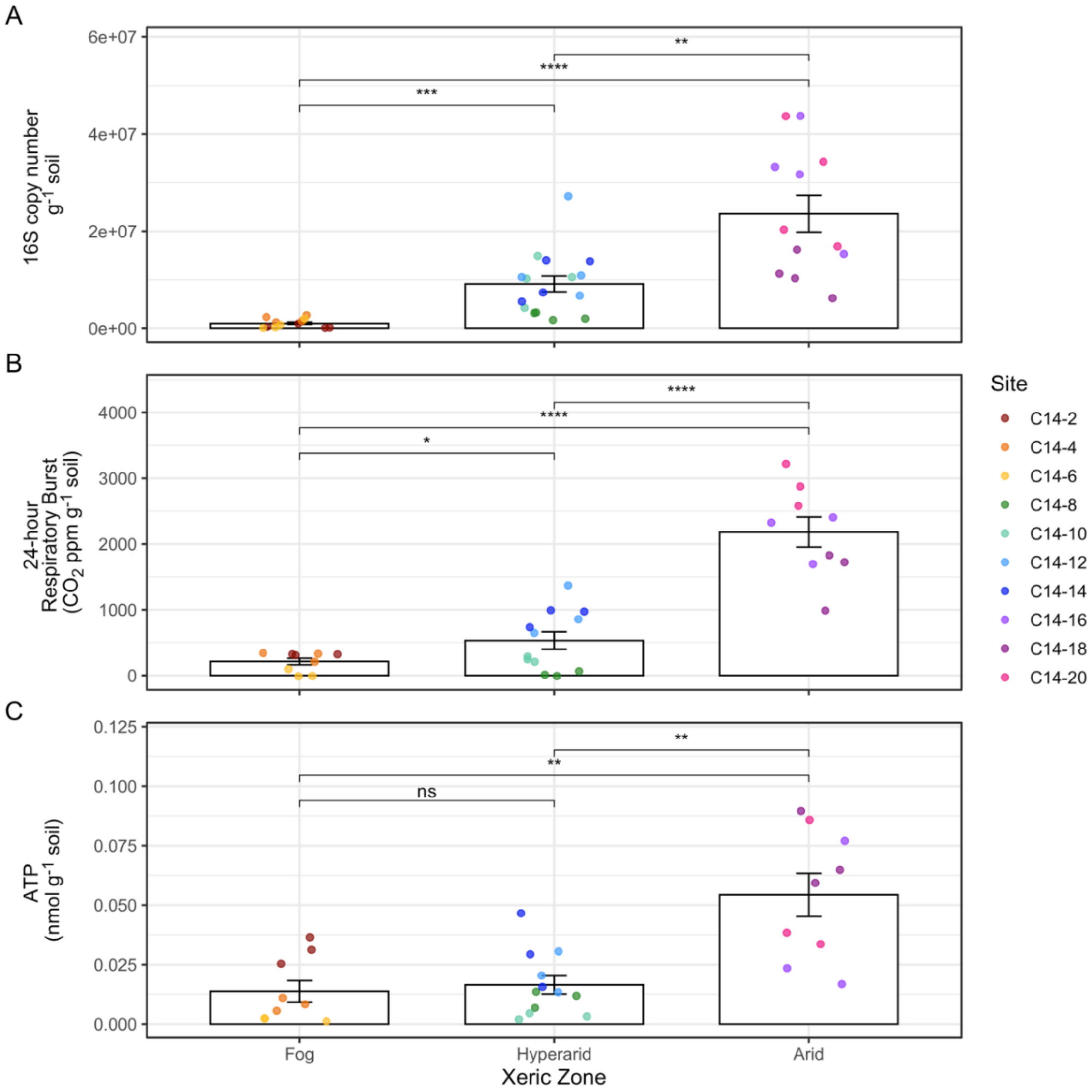


***Figure S2: Average microbial activity and abundance per xeric zone across the Namib Desert C14 transect.*** *Comparisons are made between each xeric zone, and significance levels are represented as follows:* $p \leq 0.05$ *= *,* $p \leq 0.01$ *= **,* $p \leq 0.001$ *= ***,* $p \leq 0.0001$ *= ****. Data points represent biological replicates per site (n = 4 for A, n = 3 for B and C) and error bars represent standard error. A) Total 16S rRNA gene copy numbers in sites across each xeric zone. Total copy number increased significantly between each xeric zone and was lowest in the Fog zone. B) Cumulative* $CO_2$ *respiratory burst 24 h after moisture addition. Cumulative respiratory burst increased significantly between each xeric zone and was lowest in the Fog zone. B) Intracellular ATP concentration. No significant difference in average intracellular ATP was recorded between Fog and Hyperarid zone sites.*

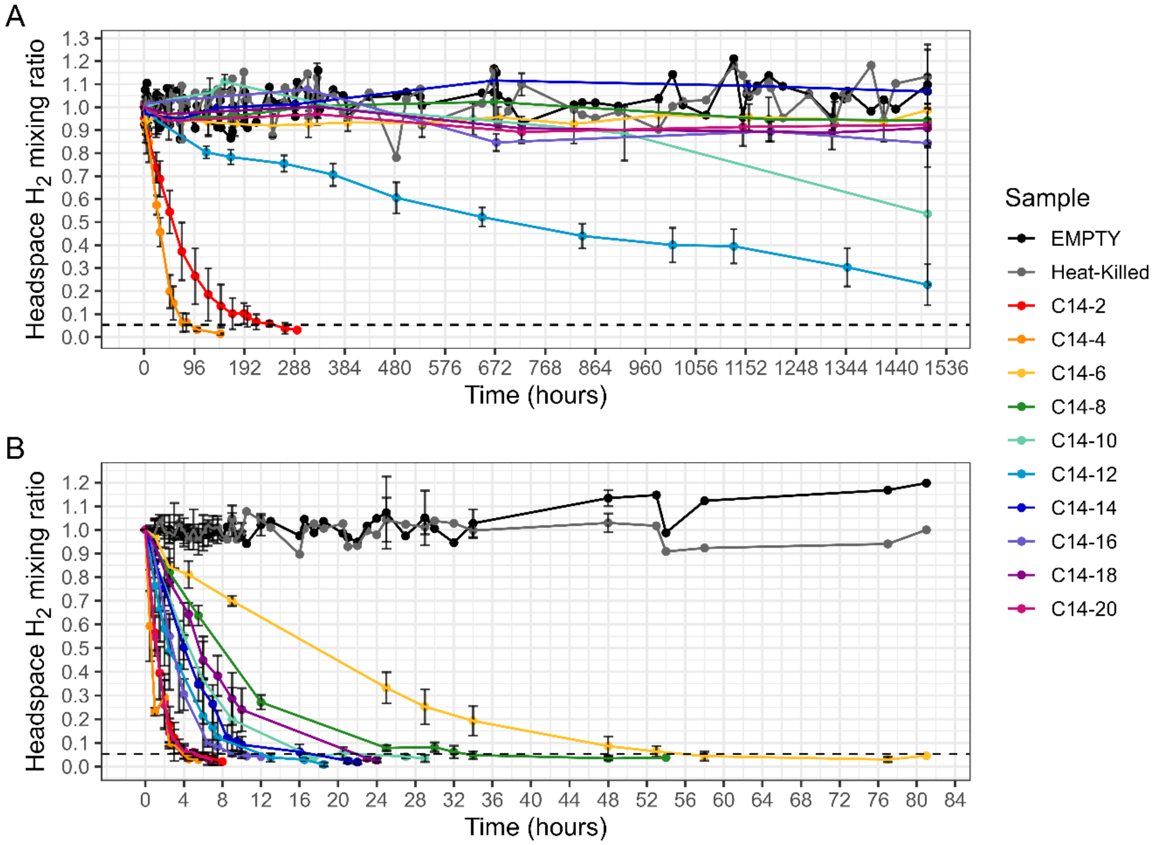


***Figure S3: Headspace hydrogen uptake by dry and wet soil Namib Desert C14 transect microcosms.*** *Data points represent the mean of five biological replicates for each site and error bars represent standard deviation. Headspace mixing values for heat-killed and empty vials from each site are represented as aggregate means across all sites for clarity. Atmospheric hydrogen concentration (530 ppbv) is represented by the dotted black line. A) Headspace hydrogen mixing ratio in dry soil 2 g microcosms incubated at 30 °C in the dark. B) Headspace hydrogen mixing ratio in wetted soil 2 g microcosms incubated at 30 °C in the dark.*

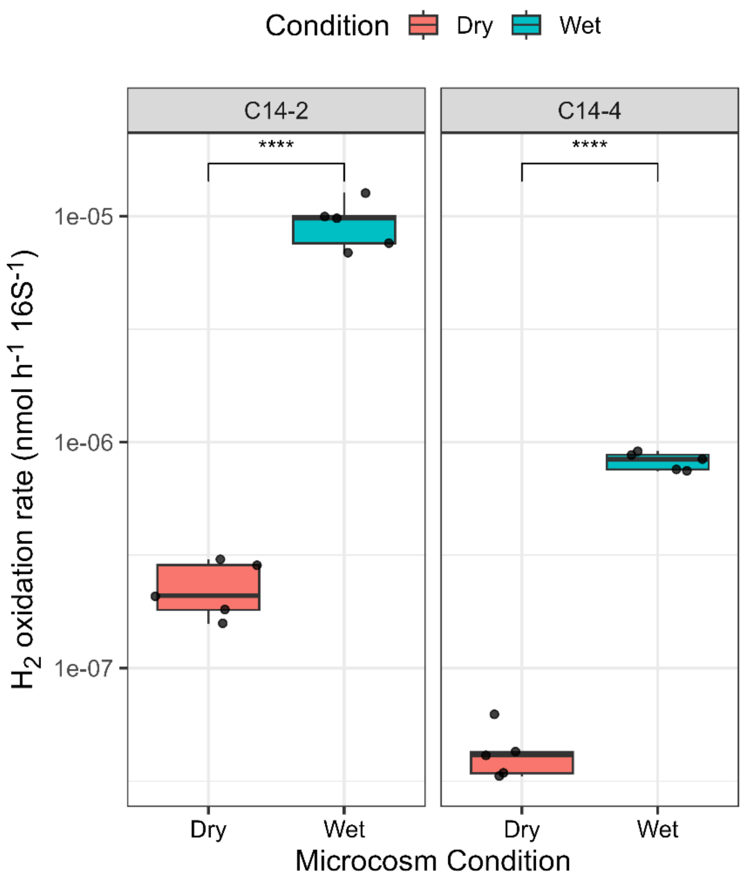


***Figure S4: Comparison of 16S-normalised atmospheric hydrogen oxidation rate in wet and dry microcosms from Namib Desert C14 transect sites C14-2 and C14-4.*** *Data points represent five biological replicates for each site and condition. Rates increased significantly in wetted conditions in both sites. Significance levels are assigned as follows:* $p \leq 0.05$ = *, $p \leq 0.01$ = **, $p \leq 0.001$ = ***, $p \leq 0.0001$ = ****.

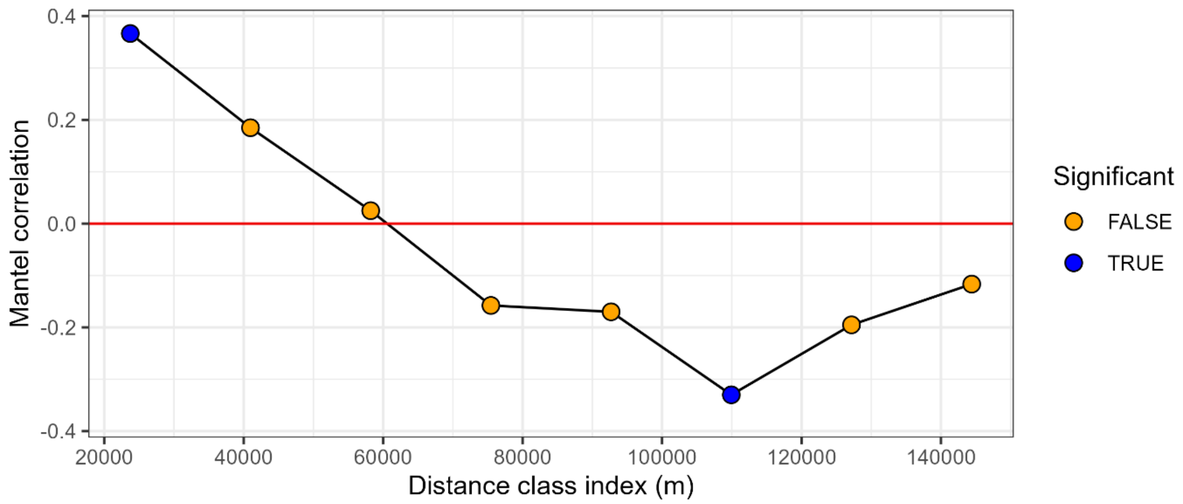


***Figure S5: Mantel correlogram showing the correlation between spatial distance and community composition based on weighted UniFrac distances.*** *Points represent distance classes calculated from GPS coordinates using the Haversine method. Significant values (p ≤ 0.05) are represented in blue, non-significant values (p > 0.05) are represented in orange.*

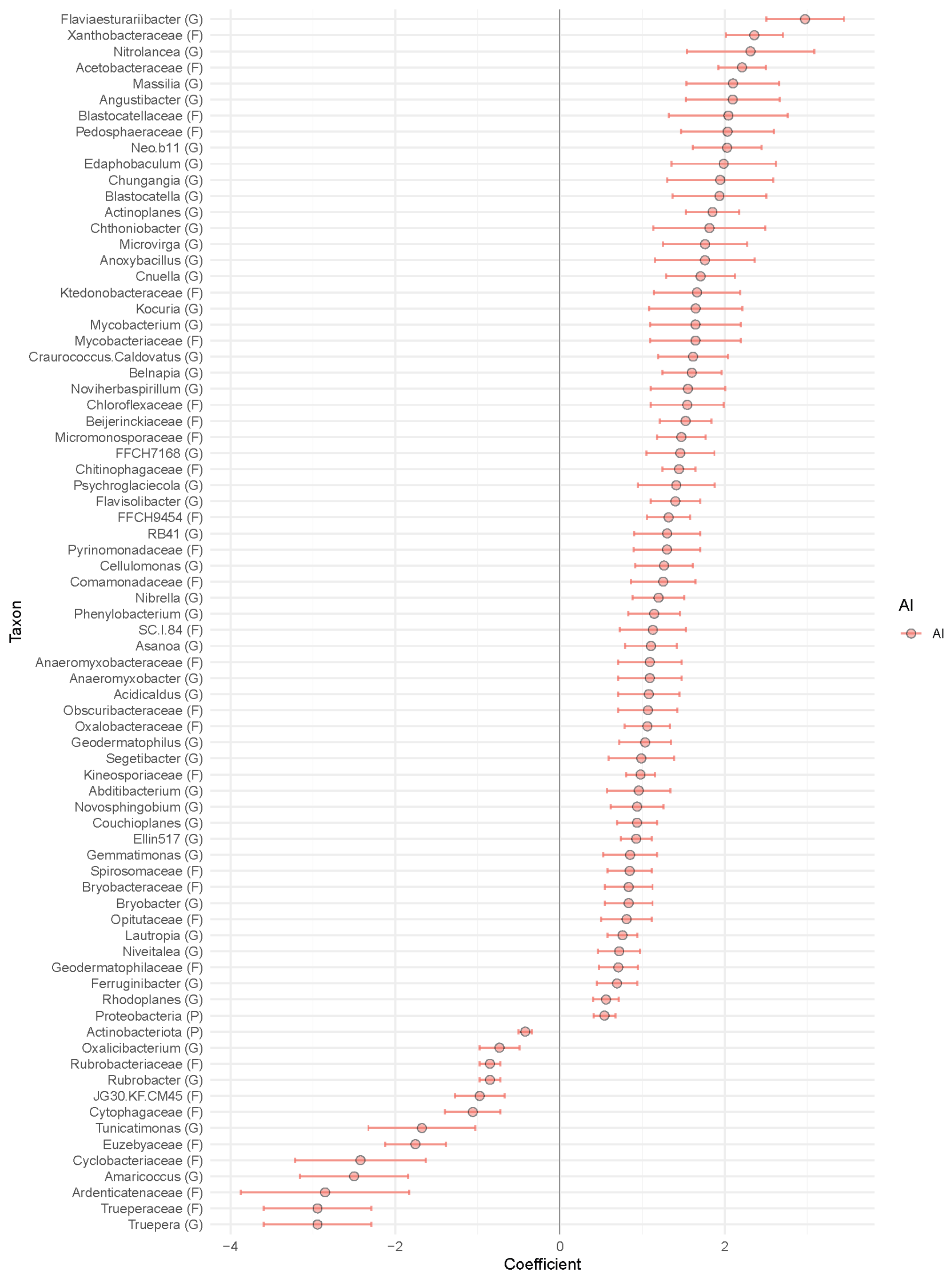


***Figure S6: Differential abundance of taxa across a Namib Desert aridity transect by Aridity Index.*** *Coefficient is a measure of the difference in relative abundance of taxa in sites according to Aridity Index (AI). Taxonomic levels are indicated as follows: (P) = Phylum, (F) = Family, (G) = Genus. Negative coefficients indicate higher relative abundance in sites with low AI. Only coefficients with adjusted q-values ≤0.20 are shown. Error bars represent standard error.*

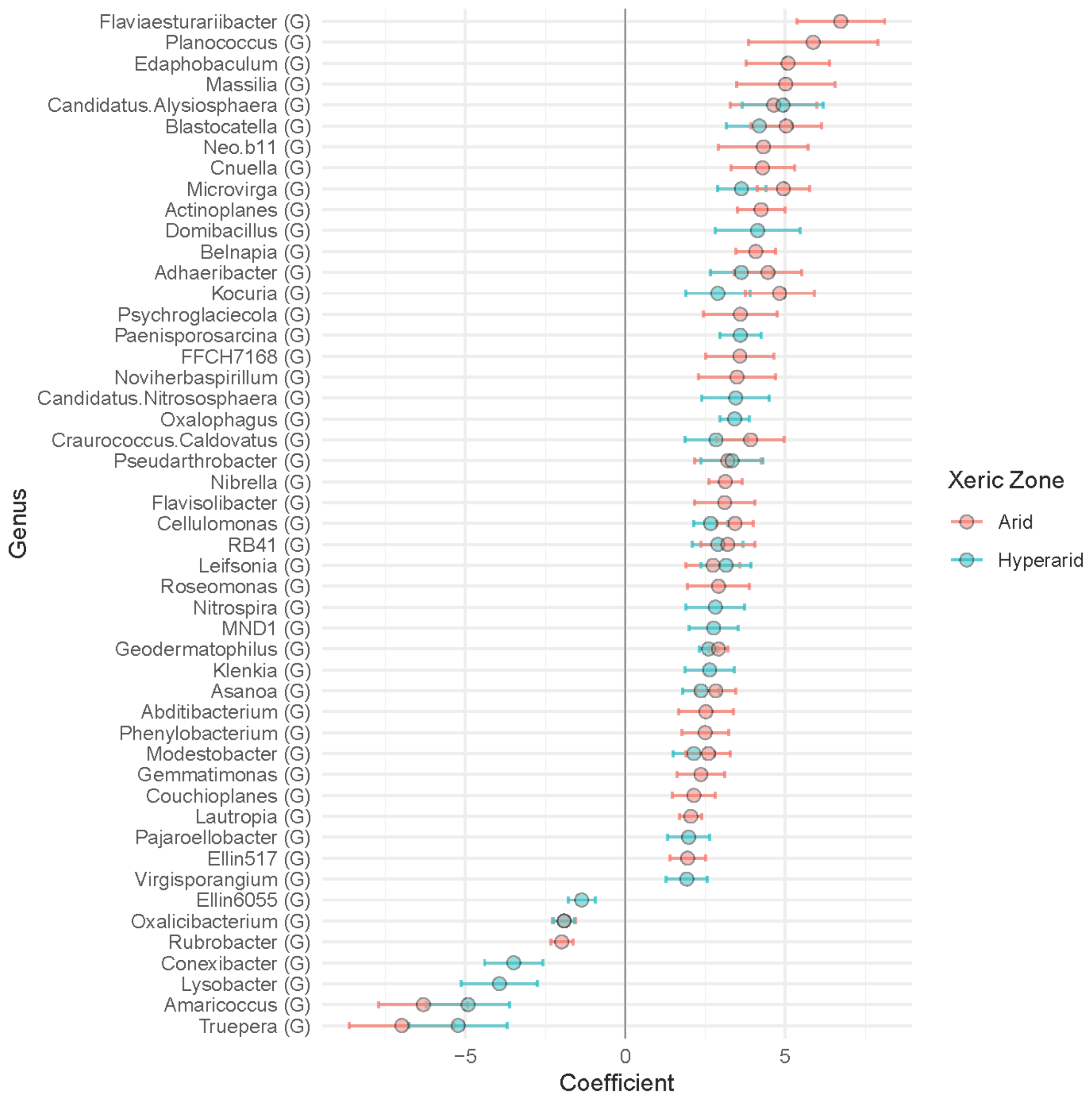


***Figure S7: Differential abundance of genera across xeric zones of a Namib Desert aridity transect, relative to the Fog zone.*** *Coefficient is a measure of the difference in relative abundance of taxa in Hyperarid and Arid zone sites, relative to Fog zone sites. Negative values indicate greater relative abundance within the Fog zone. Only coefficients with adjusted q-values ≤0.20 are shown. Error bars represent standard error.*
